# Novel Murine models of Mania and Depression

**DOI:** 10.1101/2022.03.16.484557

**Authors:** Binjie Chen, Maosheng Xia, Ming Ji, Wenliang Gong, Dianjun Zhang, Xinyu Li, Siman Wang, Yuliang Feng, Xiafang Wu, Lulu Cui, Alexei Verkhratsky, Baoman Li

## Abstract

Neuropathological mechanisms triggering manic syndrome or manic episodes in bipolar disorder remain poorly characterised, as the research progress is severely limited by the paucity of appropriate animal models. Here we developed a novel manic mice model by combining a series of chronic unpredictable rhythm disturbances (CURD), which include disruption of circadian rhythm, sleep deprivation, exposure to cone light, with subsequent interference of followed spotlight, stroboscopic illumination, high temperature stress, noise disturbance and foot shock. To validate this novel manic model, we used multiple behavioural and cell biology approaches comparing the CURD-model with healthy controls and depressed mice. The depression model was created by an exposure to an improved chronic unpredictable mild stress, which we defined as chronic unpredictable mild restraint (CUMR). A novel manic mice model induced by environmental stressors and free from genetic or pharmacological interventions will benefit research into pathological mechanisms of mania.

## Introduction

Neuropathological mechanisms of manic syndrome or manic episodes in bipolar disorder remain poorly characterised, as the research progress is severely limited by the paucity of appropriate animal models [1]. At present, the most commonly used murine mania models are based either on pharmacological treatments [2] or genetic modifications [3, 4]. In addition, a mouse model was made through sleep deprivation [5, 6]. However, none of these models faithfully reproduce the pathology. Treatments with amphetamine or cocaine interfere with the analysis of pharmacology of antimanic drugs and recruit pathological mechanisms of drug abuse. Genetically modified animal models are based on manipulations with the suspected risk genes of mania or BD that are not covering all clinical presentations, while models build through sleep deprivation display multiple and diverse outcomes including cognitive impairments or major depression [7, 8]. Previous attempts to establish manic models ignored many environmental factors (such as circadian and seasonal disturbance) that are known to contribute to the aetiology of mania [9].

## Materials and methods

### Materials

Most chemicals, including Alexa 555-conjugated ovalbumin (45 kDa) and Fluorescein 3000 MW Anionic dextran (3 kDa) were purchased from Invitrogen (CA, USA). D-Amphetamine (14204) was purchased from Cayman (Michigan, USA). Pentobarbital sodium (P3761), primary antibody of 5-HTT (SAB2502028), β-actin (A5441) and serotonin (14927) were purchased from Sigma (MO, USA). Primary antibody of NeuN (PA5-78693) was purchased from Thermo Fisher Scientific (Waltham, MA USA). 4′,6-diamidine-2-phenylindole dihydrochloride (DAPI) were purchased from Thermo Fisher Scientific (CA, USA), Lithium carbonate (sc-203109) and primary antibody of c-fos (sc-271243) were purchased from Santa Cruz Biotechnology (Dallas, Texas, USA).

### Animals

Male wild type C57BL/6 mice (#000664; aged 10-12 weeks; weight of 25-35 g) were purchased from the Jackson Laboratory (Bar Harbor, ME, USA). Animals were raised in standard housing conditions (22 ± 1[; light/dark cycle of 12/12h), with water and food available *ad libitum*. All experiments were performed in accordance with the US National Institutes of Health Guide for the Care and Use of Laboratory Animals (NIH Publication No. 8023) and its 1978 revision, and all experimental protocols were approved by the Institutional Animal Care and Use Committee of China Medical University, No. [2020]102.

### Chronic unpredictable rhythm disturbance (CURD) regimen

Male mice were exposed to the following stressors for 3 weeks. The normal rhythm was alternated 12 hours between light (7:00-19:00) and darkness (19:00-7:00). The circadian rhythm was interfered with by shortening 6 hours in the darkness time (model 1) or in the light time (model 2), sleep deprivation (6 hours), stroboscopic illumination in the dark (120/min for 12 hours), one solid cone light was turned on irregularly during the dark period (12 hours), one spotlight always followed the mice during the darkness (12 hours), environmental temperature increase to 40-45°C (30 minutes), stroboscopic illumination in the dark (120/minute for 12 hours), noise (100 dB for 12 hours), foot shock (0.8 mA for 2 seconds). Two of these nine stressors were randomly selected and applied once everyday. The detailed treatments for everyday were shown in supplementary Table 1.

### Chronic unpredictable mild restraint (CUMR) regimen

Male mice were exposed to the following stressors for 3 weeks: restricting activity (4 hours), damp bedding (12 hours), cage shaking (40/minute for 10 minutes), tail suspension (10 minutes), forced swimming (10 minutes 25°C), 45° tilting cage (12 hours). Two of these six treatments were randomly scheduled once everyday, and continued for 3 weeks, the detailed treatments for everyday were shown in supplementary Table 1.

### Chronic unpredictable mild stress (CUMS) regimen

As described previously, male mice were exposed to the following stressors for 3 weeks: water and food deprivation (12 hours), cage tilt 45° (12 hours), stroboscopic illumination in the dark (120/ minute for 12 hours), noise (120 db for 12 hours), swimming (5 minutes), tail suspension (5 minutes), damp living environment (12 hours), and cage shaking (40/minute for 5 minutes), restricting activity (4 hours) were administered, two of these treatments were randomly scheduled once everyday, and continued for 3 weeks [10].

### Ethogram

We assessed the mice status according to general appearance parameters (GAP) assessments. A score of 0 or 1 for the categories of activity, posture, breathing pattern, coat condition, and interaction with other mice were given. If the parameter was normal, it was recorded as 0, if the parameter was abnormal, it was recorded as 1. The higher the score, and the worse the state of the mice [11, 12].

### Sucrose preference test

The sucrose preference is a reward-based test and a measure of anhedonia. At first, test mice were adapted to 2.5% sucrose solution for 48 hours. In brief, after 12 hours of food and water deprivation, mice were provided with two pre-weighed bottles, including one bottle that contained 2.5% sucrose solution and a second bottle filled with water, for 12 hours. The total intake volume of pure and sucrose water were calculated, separately. The percentage preference was calculated according to the following formula: % preference = [sucrose intake/ (sucrose + water intake)] × 100% [10, 13, 14].

### Sucrose pellets preference test

The sucrose pellets preference test is additional reward-based test and a measure of anhedonia. All pellets were prepared by mixing the same weight of flour and water. The white and yellow pellets were supplemented with 0.5 g tasteless food colouring, 1 g of sucrose was randomly added to white and yellow pellets. Every pellet weigh 20 g. To avoid colour interference with the appetite, the double-blind experiments were designed, the preparation of pellets and data statistics were operated by different technicians. Before the formal test, the mice were adapted to the sweet sucrose pellets for 48 hours. Then, after 12 hours of food and water deprivation, mice were provided with sweet and unsweet of pre-weighted pellets for 6 hours. Total weight of every colour pellet remaining at the end of 6 hours was recorded. The sucrose pellets preference was calculated according to the formula % preference = [sweet pellets intake/ (sweet + unsweet pellets total intake)] × 100%.

### Tail suspension test (TST)

The tail suspension is a behavioural despair-based test. Mice were suspended by its tail around 2 cm from the tip at a height of 20 cm. Behaviour was recorded for 6 minutes. The duration of immobility in the last 4 minutes was calculated by the Labstate software (ver. 2.0) (YHTSM, Wuhan Yihong Technology Co., Ltd China) [8, 15].

### Forced swimming test (FST)

The forced swimming is a despair-based behavioural test. Each mouse was trained to swim for 15 minutes on the first day. Next day the mouse was put into a glass cylinder that contained 30 cm deep water (25 ± 1°C) for 6 minutes. The time of immobility was recorded during the last 4 minutes period which followed 2 minutes of habituation [16].

### Open filed test

The open field test evaluates the autonomous activity behaviour, exploration behaviour and anxiety of experimental animals in a new environment. The mice were placed in the open field box (50 × 50 × 50 cm), the central area covers one-third of the total open field area and behaviours were recorded for 5 minutes by the Labstate (ver. 2.0) system (YHOFM, Wuhan Yihong Technology Co., Ltd China). The parameters used for analysis included the total travel distance and time spent in the central area [17].

### Three-chambered sociability test

Three-chambered sociability test measures the social exploration of mice. The three-chamber device is a Perspex box (60 × 40 cm). The device has two gated walls dividing it into three chambers: empty, central, and <u>stranger</u>. With the gates to both chambers closed, each test mouse was placed in the central chamber for 5 minutes and then removed. A stranger mouse was placed into the transparent plastic cage in the stranger chamber to avoid direct contact with the test mice, while the cage on the other side was left empty. In the test, the gates open for the test mice to explore the whole chamber, the mice were allowed to roam freely in three chambers for 10 minutes. The movements of mice and time spent in each chamber and in social circles (the area around the strange mice cage 3 cm from the cage is defined as social circle) were recorded by a video-tracking system Labstate (ver. 2.0) (YHSAM, Wuhan Yihong Technology Co., Ltd China). Chamber duration rate (%) = chamber time/total time×100%, social circles duration rate (%) = social Circles time/total time×100% [18].

### Pentobarbital induced sleep test

Pentobarbital induced sleep test is a common method for evaluating sleep in mice. The test was set to take place between 1: 00 pm and 5: 00 pm. Each test mice was intraperitoneally injected with pentobarbital sodium (50 mg/kg). When mouse is placed in dorsal decubitus, it turns back to its normal position immediately executing the righting reflex. When the righting reflex disappears for more than 30 seconds the mouse was considered asleep. The sleep latency (the time spent between injection of pentobarbital sodium and disappearance of the righting reflex) and sleep duration (the time spent between the mice asleep and appearance of the righting reflex) were recorded [19].

### Total sleep time calculation

In order to explore the sleep duration, we conducted a 24 hours video recording, from which the sleep duration was measured manually.

### Morris water maze test

The Morris water maze test was a spatial learning and memory test. The mice were trained 5 consecutive days daily with four trials, during this period, the mice were trained from different a starting quadrant to locate and escape onto the platform. The platform position was fixed throughout the test. Animals that failed to find the location within 60 seconds were guided to the platform and were allowed to remain on it for 20 seconds. On the sixth day, the platform was removed, and the mice were given 60 seconds to explore, and the time spent in the target quadrant was collected for each mouse [10, 14].

### Rotating rod test

Rotating rod test assesses the balancing ability as a proxy for motor control by recording time spent on the rotating rod. Each mouse was placed on a rotating bar, which was set to a rotation speed of up to 20 rpm during the test. The time spent on the rotating bar was recorded as the latent period. The latency before falling was recorded using a stop watch, with a maximum of 90 seconds [10, 14, 17].

### Pole test

Each mouse was paced head-upward on the top of a vertical rough-surfaced pole (diameter 1 cm; height 55 cm). The turn downward from the top of pole (T-turn time) and the descend to the floor (T-LA) time was recorded [10, 20].

### Bite ability test

The bite ability test was used to evaluate the bite force of mice, which could indirectly reflect the irritability and physical strength of the mice. Mice were given an apple tree branch of the same shape, size and weight between 1: 00 am and 3: 00 am. The weight of the torn debris was measured.

### Monitoring glymphatic system

Mice were anaesthetised with a mixture of ketamine (100 mg/kg) and xylazine (10 mg/kg) by intraperitoneal injection. The fluorescence tracers (OA555, FITC-D3) were reconstituted in artificial cerebrospinal fluid (ACSF) at a concentration of 0.5%. Mice were anesthetised and fixed in a stereotaxic frame while the posterior atlanto-occipital membrane was surgically exposed. Using a 30 GA needle, the tracer was infused into the subarachnoid CSF via cisterna magna puncture at a rate of 2μl/min for 5 minutes (10μl total volume). 30 minutes after the start of infusion, anesthetised animals were transcardially perfusion fixed with 4% paraformaldehyde (PFA). Brain tissue was cut into 50 um slice and was imaged using Carl Zeiss Axio Scan microscope (Promenade 10, Jena, Germany) [10].

### Western blotting

Using bovine serum albumin (BSA) as the standard, the protein concentration of the sample was determined by the Lowry method. Each sample contained 100 g protein was added into 10% SDS-polyacrylamide gel electrophoresis. After electrophoretic separation and the gels were transferred to polyvinylidene fluoride (PVDF) membranes, the samples were blocked by 5% skimmed milk powder for 1 hour, and membranes were incubated overnight with the primary antibodies, specific to either 5-HTT at 1:1000 dilution, β-actin at 1:3000 dilution. After washing, specific binding was detected by horseradish peroxidase-conjugated secondary antibodies. Staining was visualised by electrochemiluminescence (ECL) detection reagents and analysed with an Electrophoresis Gel Imaging Analysis System (MF-ChemiBIS 3.2, DNR Bio-Imaging Systems, Israel). Band density was measured with Window AlphaEaseTM FC 32-bit software [21].

### Immunofluorescence (IF)

The anaesthetised mice were perfused through the heart with 4% paraformaldehyde (PFA) for 15 minutes. After dissection the brain tissue was cut into 60 μm slices. Brain slices were permeabilised by incubation for 1 hour with donkey serum. Primary antibodies against *c-fos* were used at 1:100 dilution, against NeuN was used at 1:100 dilution. And nuclei were stained with marker 4’, 6’-diamidino-2-phenylindole (DAPI) at 1:1000 dilution. The incubation with the primary antibodies were overnight at 4 °C and then donkey anti-mouse or anti-rabbit Alexa Fluor 488/555 conjugated secondary antibodies were incubated for 2 hours at room temperature. Images were captured using a confocal scanning microscope (DMi8, Leica, Wetzlar, Germany) [22].

### Microdialysis and HPLC-MS analysis

Mice were anaesthetised by a mixture of ketamine (100 mg/kg) and xylazine (10 mg/kg) by intraperitoneal injection. A guide cannula (CMA 7, CMA Microdialysis, Stockholm, Sweden) was implanted into the right prefrontal cortex (coordinates: anteroposterior 1.75 mm, mediolateral 0.75 mm, dorsoventral 1.5 mm). A microdialysis probe (CMA 7; molecular weight cut-off, 6,000 Da) was inserted through the cannula 24 hours before the start of experiments. Artificial cerebrospinal fluid (ACSF) was perfused through the microdialysis probe at 1 *μ*L/min, and samples were collected 3 hours after probe insertion. The interstitial fluid 5-HT levels were measured immediately [8]. Analyses of serotonin were performed with a high-performance liquid chromatography (HPLC; Agilent 1260 Infinity LC system) tandem with a mass spectrometry (MS) system, Agilent 6420 triple-quad mass spectrometer (Agilent Technologies, CA, USA). A mobile phase composed of acetonitrile (ACN), solvent A, and 0.2% formic acid in water, solvent B, was used. In the HPLC-MS system, 5 *μ*L of the samples was injected into a 120 SB-C18 column (Poroshell, Agilent, 46×100 mm). The experiments were effectuated at 20°C (room temperature) for 5 min with an elution gradient (0-2 min: 0.25 mL/min; 2-5 min: 0.5 mL/min; 0-5 min 70% solvent A, 30% solvent B). Before injections, the column was equilibrated for 15 min. The mass spectrometer parameters were as follows: positive multiple reaction monitoring mode MRM (177.2→160.2; 177.2→132.2), positive electrospray ionisation (ESI), collision energy: 15, 10 L/min heating gas flow, 300°C interface temperature, 3 L/min nebulising gas flow, 300 °C DL temperature, 400°C heat block temperature, and 10 L/min drying gas flow. The detection limit was 1 pmol per injection [23]. The quantification was performed using peak area ratios from calibration standard curves and was normalised to the total protein levels in the samples as determined by the Lowry method.

## Results

We used a series of environmental interventions to produce a novel murine mania model. We defined the complex of these interventions as a chronic unpredictable rhythm disturbance (CURD), which subjected animals for a regimen of eight stressors applied for three weeks (Fig. 1a-p). As shown in Fig. 1a, after one week of the 12/12 hours light/dark environmental adaptation, the mice were treated by two out of eight randomly selected stressors everyday for three weeks. Detailed treatments protocol is shown in Table 1, while stressors are summarised in Fig. 1b-1i.

**Figure 1.**
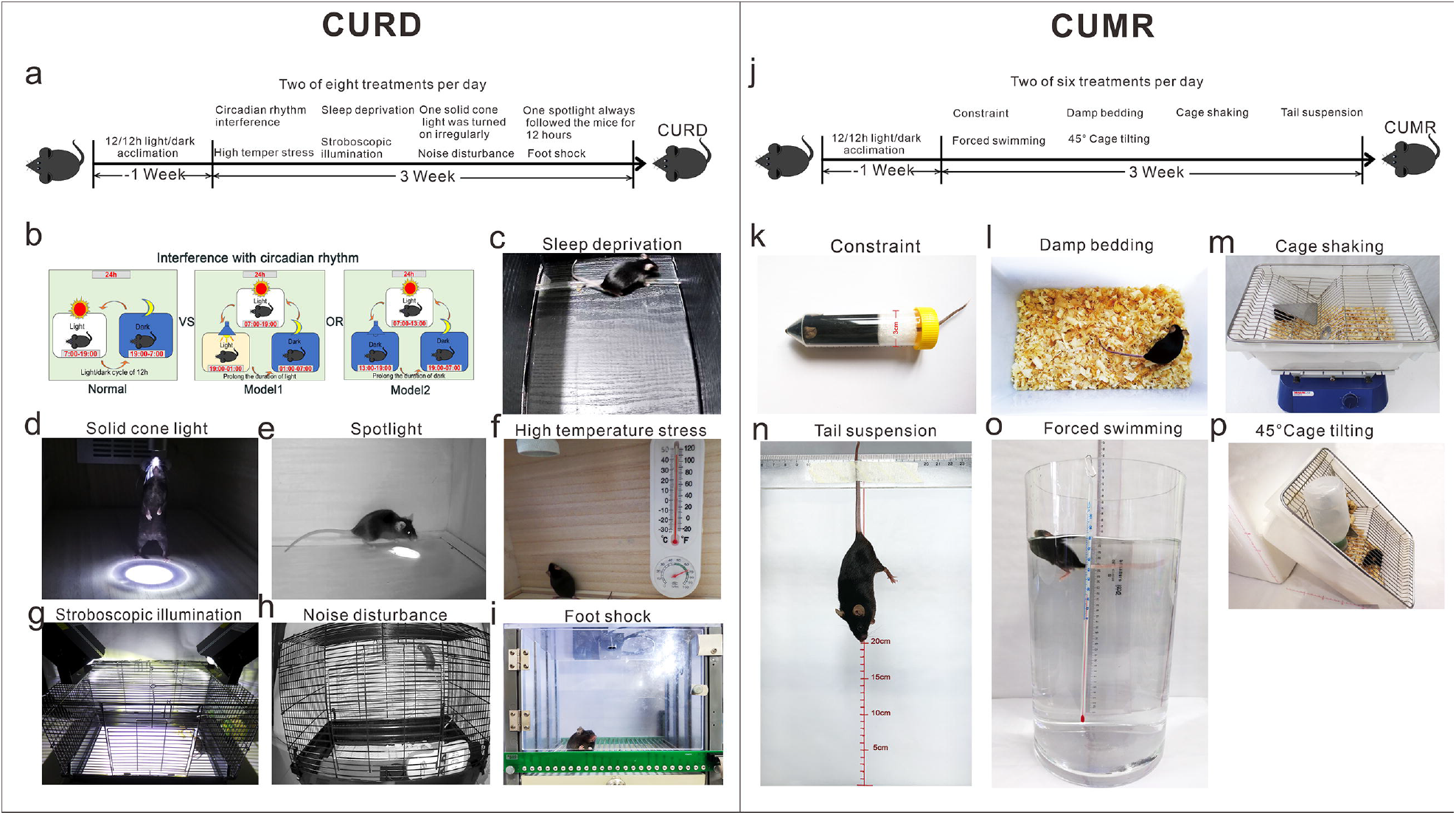
The protocol and stressors used for chronic unpredictable rhythm disturbance (CURD) and chronic unpredictable mild restraint (CUMR). To establish CURD model, the interference of circadian rhythm (b), sleep deprivation (c), the interference of cone light (d), the interference of followed spotlight (e), high temperature stress (f), stroboscopic illumination (g), noise disturbance (h) and foot shock (i) were employed. Two out of these nine stressors were randomly chosen and applied every day, the whole period of disturbed rhythm was 3 weeks. To establish CUMR model, constraint (k), damp bedding (l), cage shaking (m), tail suspension (n), forced swimming (o) and cage tilting (p) were used. Two out of these six behavioral constraints were randomly selected and applied every day for 3 weeks.

The stressors employed were as follows. (i,ii) The normal rhythm was interfered as mode 1 (the prolonged duration of light) or 2 (the prolonged duration of dark) (Fig. 1b). (iii) Sleep deprivation was continued for 6 hours, beginning at 7 a.m. and ending at 1 p.m. (Fig. 1c). (iv) During darkness period a solid cone light was turned on irregularly, (Fig. 1d). (v) A spotlight was following the mouse for 12 hours (Fig. 1e). (vi) Environmental temperature was increased to 40 - 45°C for 30 minutes (Fig. 1f). (vii) Animals were exposed to two strobe lamps at 120/min for 12 hours (Fig. 1g). (viii) White noise was applied at 100 dB for 12 hours (Fig. 1h). (ix) Foot shock was set at 0.8 mA for 2 s and was repeated three times during light period (Fig. 1i). The above manipulations were employed to disturb the rhythms, including living with stressors (iv, v, vii, viii and ix), circadian stressors (i and ii), sleep by stressor (iii) and body temperature by stressor (vi).

We also developed a protocol to specifically induce depression by employing the set of restrain-related stressors. We defined this regimen as chronic unpredictable mild restraint (CUMR), which is based on and further advances the widely used chronic unpredictable mild stress (CUMS) protocol [13, 14, 24]. Major difference between CUMR and CUMS is the exclusion of rhythm disturbances such as food deprivation, group housing and stroboscopic illumination. The CUMR protocol is shown in Fig. 1j. After one week of the environmental adaptation, mice were randomly treated by two out of six randomly selected stressors every day for three weeks. Detailed treatment protocol is shown in Table 1, while stressors are summarised in Fig. 1k-1p. Six stressors include (i) behavioural constraint for 4 hours (Fig. 1k), (ii) damp bedding for 12 hours (Fig. 1l), (iii) cage shaking at 40/minute for 10 minutes (Fig. 1m), (iv) tail suspension for 10 minutes (Fig. 1n), (v) swimming in 25°C water for 10 minutes (Fig. 1o) and (vi) tilting cage at 45° for 12 hours (Fig. 1p).

Exposure to CURD and CUMR regimens resulted in development of two distinct phenotypes. Most overt was the general appearance of animals: mice subjected to CURD were hyperactive and restless, whereas mice subjected to CUMR were passive and idle (Fig. 2a and Supplementary videos 1-3). The general appearance parameters (GAP) scores were used to obtain the ethogram (every animal was given a score of 0 or 1 for the categories of activity, posture, breathing pattern, coat condition, and interaction with other mice, Fig. 2b). The GAP scores increased significantly in CURD (p < 0.0001, n = 12) and were even higher in CUMR groups (p = 0.0011, n = 12 when compared to CURD). Sucrose uptake in CURD and CUMR groups decreased significantly (Fig. 2c) to 70.47 ± 8.92% and 54.31 ± 7.71% of control group (p < 0.0001, n = 12), with sucrose preference in CURD group being higher than in CUMR group (p < 0.0001, n = 12). The total intake volume of water in CURD group increased to 131.40 ± 11.63 % of control group (p < 0.0001, n = 12; Fig. 2d), whereas in CUMR group it decreased to 43.64 ± 10.11% of control group (p < 0.0001, n = 12; Fig. 2d). To further assess the preference for solid food, we used a sucrose pellets preference test (Fig. 2e-2g). The intake percentage of sucrose sweet pellets decreased in both CURD and CUMR groups, with ratios of 80.74 ± 3.94% (p < 0.0001, n =12) and 70.47 ± 4.74% (p < 0.0001, n = 12) of control (Fig. 2e,f). The total intake weight of pellets in CURD group increased to 145.50 ± 14.18% of control group (p< 0.0001, n = 12; Fig. 2g), while in CUMR group it decreased to 47.47 ± 10.19% of the control (p < 0.0001, n = 12; Fig. 2g). Immobility times in despair related tail suspension and forced swimming tests in CURD group were significantly decreased (p<0.0001, n = 12); to the contrary the immobility time in CUMR group increased (p<0.0001, n = 12) (Fig. 2h and 2i).

**Figure 2.**
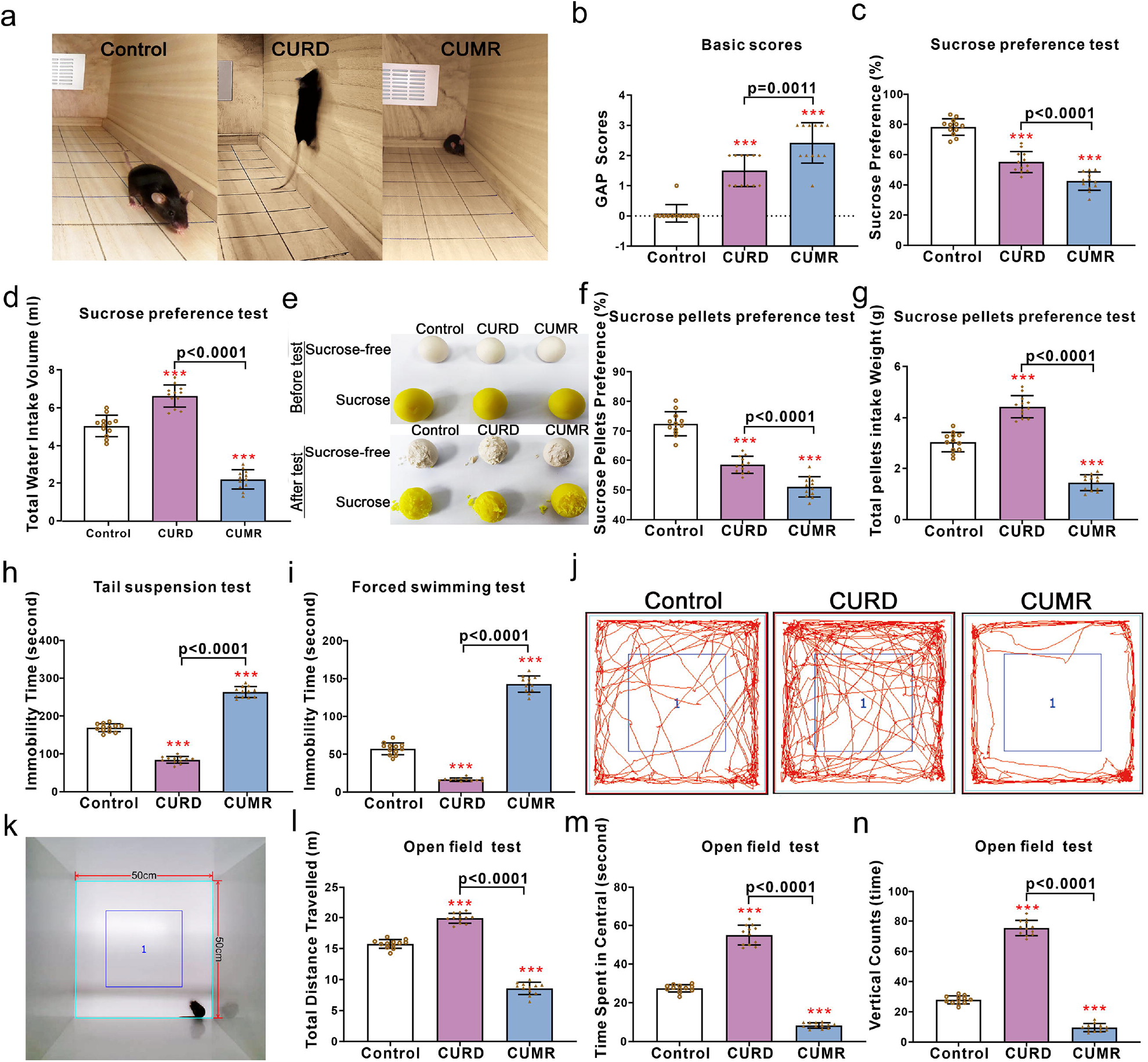
Anhedonia and anxiety behavioural measures of CURD and CUMR mice. After treatments with sham, CURD or CUMR for 3 weeks, the mice behaviours in three groups were tested. (a) The representative pictures of mice in control, CURD and CUMR groups. (b) The general appearance parameters (GAP) scores were used to assess ethogram. (c) The percentage of sucrose preference was used to test anhedonia. (d) The total intake volume of water used to test appetite. (e) The representative pictures of pellets in sucrose pellets preference test. (f) The percentage of sucrose pellets preference was operated to assess anhedonia. (g) The total intake weight of pellets was used to assess appetite. The immobility time in tail suspension (h) and forced swimming tests (i) were recorded in the last 4 minutes of 6 minutes. (j-n) In open field test, the picture of open field was shown in k. The representative trails of mice treated in control, CURD and CUMR groups were shown in j, the total travelled distance (l), the time spent in central space (m) and the times of vertical counts (n) were recorded in 5 minutes. Data are presented as mean ± SD, n = 12 per group. One-way ANOVA for comparisons including more than two groups; unpaired two-tailed t-test for two group comparisons. Significant different from control group: *p<0.05, **p<0.01, ***p<0.001.

The recorded total travel distance (anxiety related open field test, Fig 2j-2l) in CURD group increased significantly to 126.30 ± 5.04% of the control (p < 0.0001, n = 12). In contrast, the travel distance in CUMR group decreased to 54.48 ± 6.27% of control values (p < 0.0001, n = 12). Time spent in central area as well as times of vertical counts were increased in CURD group, but reduced in CUMR group (Fig. 2m and 2n; p < 0.0001, n = 12). In the three chambered sociability test (Fig. 3a, 3b), the trail of the test mouse were recorded in 10 minutes, and the representative hot points images of the test mouse in three chambers were detected for control, CUMR and CURD groups (Fig. 3c).

**Figure 3.**
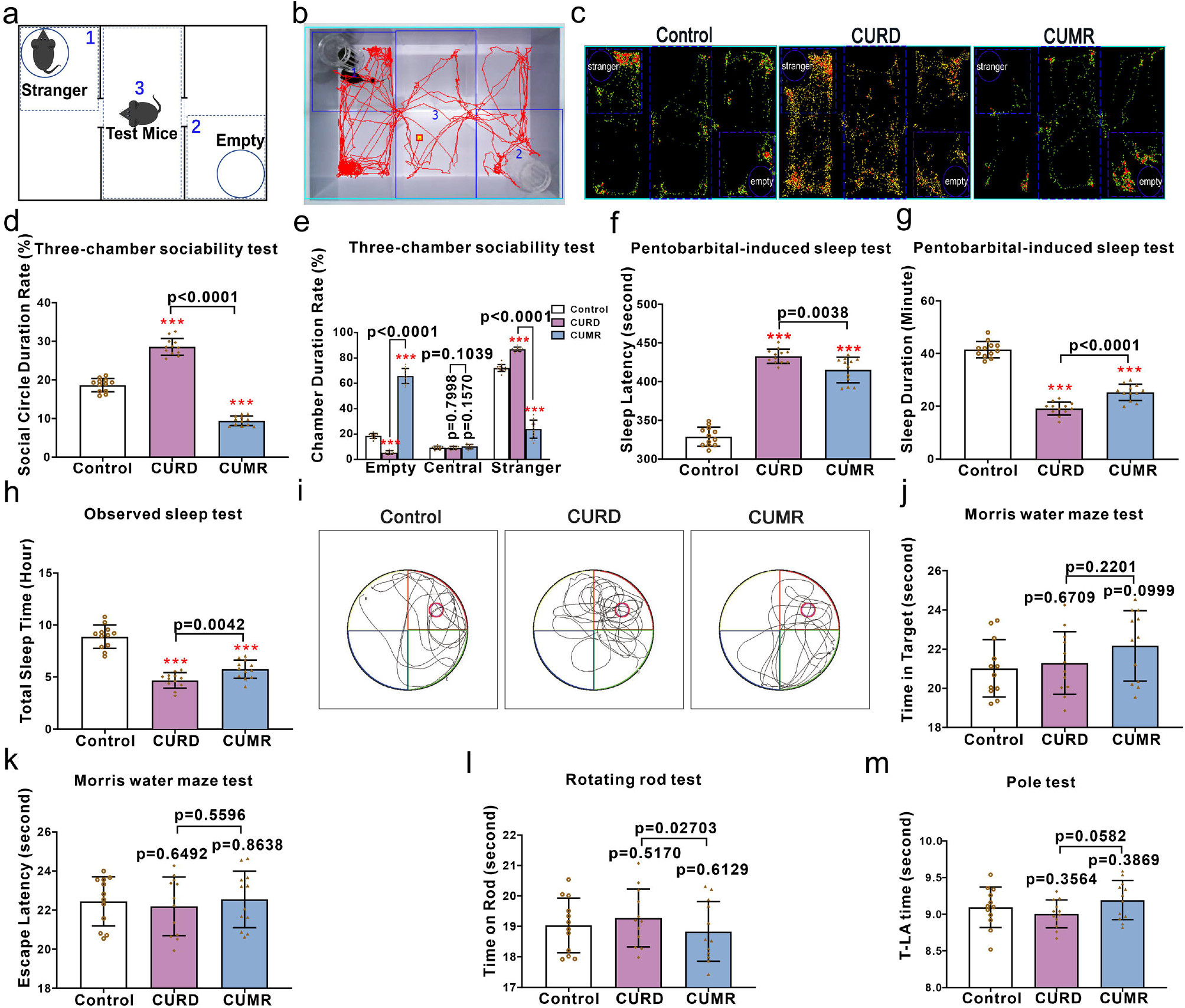
The social and cognitive behavioural measures of CURD and CUMR mice. After treatments with sham, CURD or CUMR for 3 weeks, mice behaviours were tested. (a-e) Schematic diagram of three chamber test and the representative trail of CURD treated mice are shown in a and b. The representative hot points images of the mice in control, CURD and CUMR groups (c), the ratio of social circle duration (d) and chamber duration (e) recorded for 10 minutes. After intraperitoneal injection of pentobarbital at 50 mg/kg dose, the time of sleep latency (f) and sleep duration (g) were recorded. The total sleep time of mice without interference were recorded in 24 hours (h). (i-k) In Morris water maze test, the representative trails of mice in control, CURD and CUMR groups are shown in i, the time in target quadrant (j) and the time of escape latency (k) were recorded in 60 seconds. The time on rotating rod (l) and the T-LA time in pole test (m) were recorded for control, CURD and CUMR groups. Data are presented as mean ± SD, n=12 per group. One-way ANOVA for comparisons including more than two groups; unpaired two-tailed t-test for two group comparisons. Significant different from control group: *p<0.05, **p<0.01, ***p<0.001.

Social duration ratio in CURD mice with the stranger mice increased to 153.10 ± 11.71% of control group (p < 0.0001, n = 12), while the social duration ratio of CUMR mice decreased to 50.61 ± 6.72% of control group (p < 0.0001, n = 12, Fig. 3d). As compared with control group, the time spent in the empty chamber significantly decreased in CURD group (p < 0.0001, n = 12) while being increased in CUMR group (p < 0.0001, n = 12). The time spent in stranger chamber increased in CURD group (p < 0.0001, n = 12) and decreased in CUMR group (p < 0.0001, n = 12). The residence time in central chamber was the same in all groups (Fig. 3e). In pentobarbital-induced sleep induction test (an intraperitoneal injection of 50 mg/kg of pentobarbital), the sleep latency times were prolonged in both CURD and CUMR mice (p < 0.0001, n = 12; Fig. 3f). In contrast, the sleep duration time of CURD and CUMR groups was significantly reduced (p < 0.0001, n = 12; Fig. 3g). The total sleep time (recorded for 24 hours without any interference) of CURD and CUMR mice was less than in control (p < 0.0001, n = 12; Fig. 3h). The total sleep time of CURD group was shorter than CUMR treated mice (p = 0.0042, n = 12; Fig. 3h). In Morris water maze test (Fig. 3i), the time spent in target quadrant and the time of escape latency were similar in all groups (Fig. 3j and 3k). Likewise, the motor function (assessed by rotating rod and pole tests), was not affected. Neither time kept on the rod nor the T-LA time (turn downward from the top of pole, the T-turn time, and descend to the floor) differed between groups (Fig. 3l and 3m).

We further compared the behavioural performance of mice exposed to CURD or CUMR regimen with pharmacological mania mice models (Fig. 4). The amphetamine (AMP) model [25, 26] was established by intraperitoneal injection of 2.5 mg/kg amphetamine for 10 days. For the corticosterone (COR) model 20 mg/kg corticosterone was intraperitoneally injected for 28 days [15, 27]. Sucrose uptake percentage significantly decreased in CURD, AMP, CUMR and COR groups (p < 0.0001, n = 20; Fig. 4a). The sucrose intake in CURD group was lower than in AMP group (p = 0.0053, n = 20; Fig. 4a), while it was higher in CUMR group when compared with COR group (p = 0.0041, n = 20; Fig. 4a). The total intake volume of water (Fig. 4b), of CURD and AMP groups were significantly increased (p<0.0001, n=20) when compared with control group, with CUMR and COR groups being decreased much more (p < 0.0001, n =2 0). The intake of solid sweet sucrose pellets was significantly decreased in CURD, AMP, CUMR and COR groups as compared with control group (p < 0.0001, n = 20; Fig. 4c). When compared to the control group, the total intake weight of pellets was significantly increased in CURD and AMP groups (p < 0.0001, n = 20; Fig. 4d), but decreased in CUMR and COR groups (p < 0.0001, n = 20; Fig. 4d). In the despair related tail suspension and forced swimming tests (Fig. 4e and 4f), the immobility time in CURD and AMP groups were all decreased significantly as compared with control group (p < 0.0001, n = 20). The immobility time of CURD group was much higher when compared with AMP treated mice in tail suspension test (p= 0.0004, n = 20) and in forced swimming test (p = 0.0316, n = 20). However, in these two despair-related tests, the immobility time in both CUMR and COR groups was increased when compared with control group (p < 0.0001, n = 20). The immobility time of CUMR treated mice were significantly lower than of COR treated mice in tail suspension test (p = 0.0009, n = 20) and in forced swimming test (p= 0.0003, n = 20), respectively (Fig. 4e and 4f).

**Figure 4.**
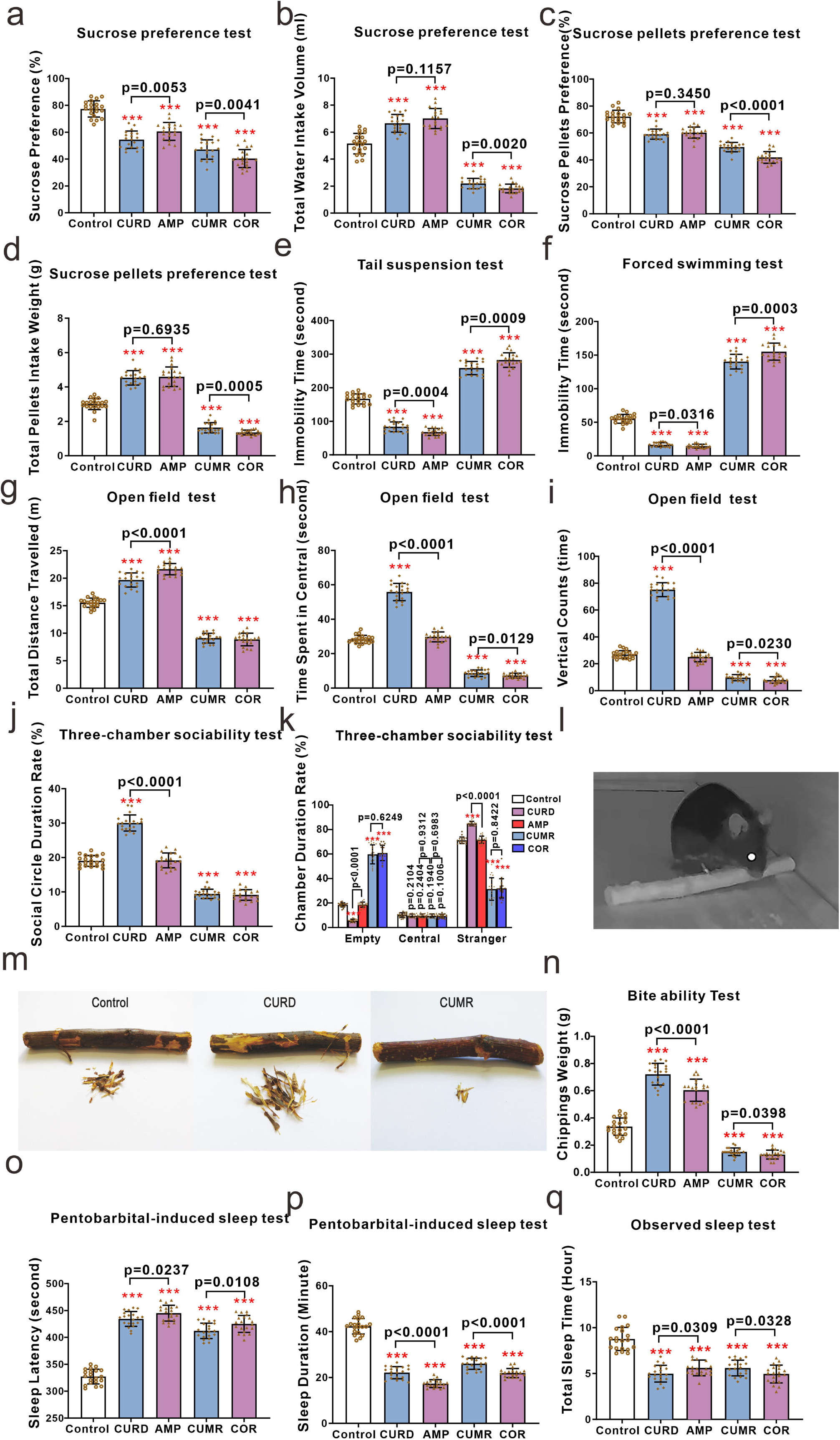
Behavioural indicators for CURD and CUMR mice as compared with amphetamine and corticosterone models. (a) The percentage of sucrose preference. (b) The total intake volume of water used to test appetite. (c) The percentage of sucrose pellets preference was operated to assess anhedonia. (d) The total intake weight of pellets was used to assess appetite. The immobility time in tail suspension (e) and forced swimming tests were recorded in the last 4 minutes of 6 minutes. (f) In the open field tests, the total travelled distance (g), the time spent in central space (h) and the times of vertical counts (i) were recorded in 5 minutes. In three-chambered sociability test, the ratio of social circle duration (j) and chamber duration (k) were recorded in 10 minutes. (l and m) The picture mice of gnawing in the darkness and the representative images of stick scrap gnawed by mice in control, CUMR and CURD groups. (n) The chippings weights gnawed by mice in 2 hours. After intraperitoneally injection of pentobarbital at 50 mg/kg dose, the time of sleep latency (o) and sleep duration (p) were recorded. The total sleep time of mice without interference were recorded in 24 hours (q). Data are presented as mean ± SD, n=20 per group. One-way ANOVA for comparisons including more than two groups; unpaired two-tailed t-test for two group comparisons. Significant different from control group: *p<0.05, **p<0.01, ***p<0.001.

Total travelled distance (open field test (Fig. 4g-4i)) was significantly increased in CURD and AMP groups (p < 0.0001, n = 20), while decreasing in CUMR and COR groups (p < 0.0001, n = 20). Travelled distance in CURD group was significantly higher than in AMP group (p < 0.0001, n = 20), while there was no significant difference between CUMR and COR groups (Fig. 4g). In the same open field test, the time recorded in the central region increased only in CURD group (p < 0.0001, n = 20), with no significant difference between control and AMP groups. The time in the central region was reduced in CUMR and COR mice (p < 0.0001, n = 20), with significant difference between CUMR and COR groups (p = 0.0129, n = 20). Calculated vertical counts in the CURD group was higher than in the control group (p < 0.0001, n = 20), whereas the vertical times were reduced in CUMR and COR mice as compared with the control group (p < 0.0001, n = 20) with no significant difference between control and AMP group (Fig. 4i). The ratio of duration time in the empty chamber (the social related three chamber test) decreased significantly in CURD group while increasing in CUMR and COR groups, with no difference in AMP group (Fig. 4j). Social circle duration ratio was significantly increased in CURD treated mice (p < 0.0001, n = 20) while decreasing in CUMR and COR treated mice (p < 0.0001, n = 20); Again, there was no difference between control and AMP groups (Fig. 4k).

Rodents have a propensity to gnaw [28]. We weighed the gnawed scrap of stick during darkness to determine the biting preference of mice (Fig. 4l). The representative images of stick scrap gnawed by mice from different groups in 120 minutes are shown in Fig. 4m. As compared with control group, the weight of scrap was significantly increased in CURD and AMP groups (p < 0.0001, n = 20), but was reduced in CUMR and COR groups (p < 0.0001, n = 20; Fig. 4n). In the pentobarbital induced sleep test, the time of sleep latency in CURD, AMP, CUMR and COR groups were all significantly higher than in control group (p < 0.0001, n = 20; Fig. 4o), whereas the sleep duration time in these four groups was lower than in control (p < 0.0001, n = 20; Fig. 4p). Similarly, the total sleep time without any interference was shortened in all four experimental groups (p < 0.0001, n = 20; Fig. 4q).

Previously we demonstrated impairment of the glymphatic system in the CUMS mouse depression model [10, 14]. We therefore assessed glymphatic system in CURD and CUMR mice, as shown in Figs. 5a and b. We monitored the distribution of fluorescence tracers OA555 (45 kDa) and FITC-D3 (3 kDa) after injecting into the posterior cistern. Fluorescence intensities of OA555 and FITC-D3 were both significantly reduced in CURD and CUMR treated mice as compared with control group. At the same time, the tracers intensities in CURD group were significantly higher than in CUMR group (n = 12). Protein expression of *c-fos* in cortical neurones may indirectly report neuronal activity [29]. Expression of *c-fos* in neurones was increased in CURD and AMP groups (p < 0.0001, n = 12; Fig. 5d), while decreasing in CUMR and COR groups (p<0.0001, n=12; Fig. 4d). Serotonin transporter 5-HTT (SERT/SLC6a4) responsible for 5-HT clearance is linked to depressive or manic symptoms [30-32]. Protein expression of 5-HTT decreased in CURD and AMP groups (p < 0.0001, n = 12; Fig. 5f), while being increased in CUMR and COR groups (p< 0.0001, n = 12). We also found significant differences between CURD and AMP groups (p = 0.0010, n = 12) and between CUMR and COR groups (p = 0.0028, n = 12) (Fig. 5f). Extracellular 5-HT measured by microdialysis increased in CURD and AMP groups (p < 0.0001, n = 12) and decreased in CUMR and COR groups (p < 0.0001, n = 12, Fig. 5g).

**Figure 5.**
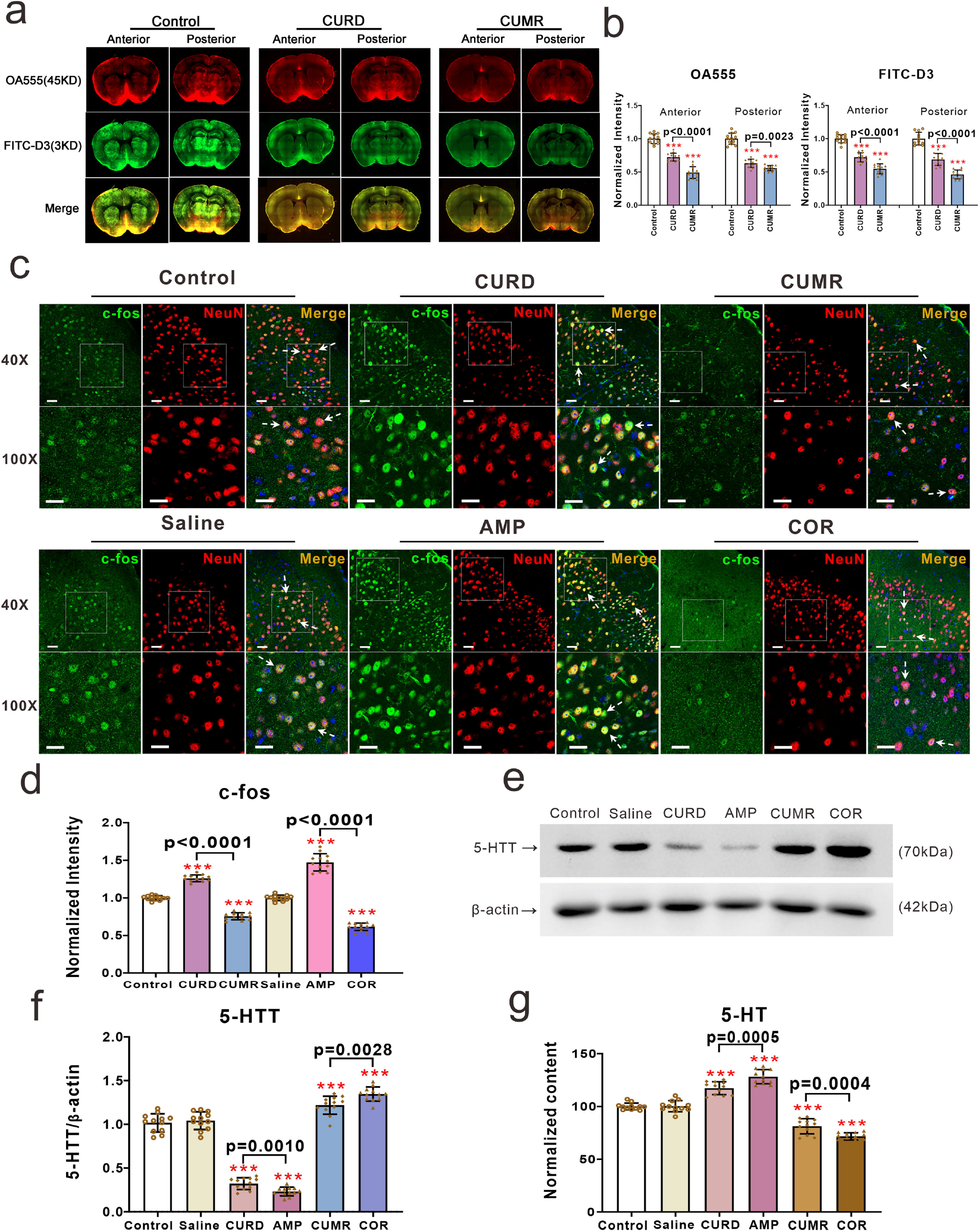
Biological indicators in the mice model of CURD or CUMR. (a-b) a large molecular weight tracer (OA555, 45 kDa) and small molecular weight tracer (FITC-D3, 3 kDa) were intracisternally injected. (a-b) Thirty minutes after injection, the animals were perfusion fixed and the whole-slice fluorescence was calculated. (a) The representative images indicate the CSF tracer penetration into the brain, OA555 (red) and FITC-D3 (green). (b) Fluorescence intensities of OA555 and FITC-D3 normalised to the intensity of the control group. (c) The representative images of fluorescence in the frontal cortex, *c-fos* (green), NeuN (red) and DAPI (blue). (d) The intensities of *c-fos* normalised to the intensity of the control group. (e) The representative protein bands of 5-HTT and *β*-actin, and the normalised intensities of 5-HTT by *β*-actin were shown in f. (g) The percentage of the extracellular 5-HT level normalised by control group in frontal cortex. Data are presented as mean ± SD, n=12 per group. One-way ANOVA for comparisons including more than two groups; unpaired two-tailed t-test for two group comparisons. Significant different from control group: *p<0.05, **p<0.01, ***p<0.001.

## Discussion

We designed new protocol for chronic stress exposures to produce divergent phenotypes of mice replicating maniac and depressive syndromes. Mice exposed to the CURD regimen present signs of manic behaviour. These animals are hyperactive in the home cages (Fig. 2a) and in the open field (Fig. 2j-2l), they exhibit reduced immobility time in tail suspension and forced swimming tests (Fig. 2h and 2i), and their exploratory behaviours (Fig. 2n) as well as social interactions (Fig. 3ak-3e) are increased. These signs were accompanied with disturbed rhythm of sleep (Fig. 3f and 3g) and shortened sleep duration (Fig. 3h). Similar behavioural manifestations, as well as sleep disturbances and increased social presence are observed in humans during manic episodes [33-35]. Some, but not all, of these mania-like behaviours including reduced immobility time, hyperactivity in cages and abnormal circadian rhythms have been reported for SHANK3 overexpression transgenic mice, considered to be a maniac model [36]. The anhedonia related sucrose preference (both in water and in pellets) is decreased in CURD group. At the same time, the total intake water volume and pellets consumption were increased, indicates that manic mice are appetite hyperactive, without distinguishing food for its taste (Fig. 4a-4d). In almost all tests, the behavioural performance observed in CURD treated mice was similar to AMP pharmacological induced mice, whereas the CUMR restraint mice and COR mice also displayed similar behavioural representation (Fig. 4). Besides establishing an effective behavioural manic model, we produced an improved depression model using newly developed CUMR protocol. Mice espoused to CUMR protocol display depressive-like behaviours. In contrast to CURD mice, animals in CUMR and COR groups showed reduced social and exploratory performance.

The CURD mania model is more comprehensive than the pharmacological (amphetamine) model. It is well documented that in healthy humans, 20 mg amphetamine increase motor activity without increasing specific exploration and without marked effects on spatial patterns of activity [37]. The amphetamine induced behaviours are inconsistent with the performance observed in either the manic or non-manic phases of bipolar disorder, which is characterised by both hyperactivity and increased exploration [38-41]. Similarly, animals acutely injected with amphetamine to model mania present hyperactivity alone with no effect on exploratory behaviour [42].

Nearly all patients suffering from mood disorders have significant disruptions in circadian rhythms and the sleep/wake cycle [43], with altered sleep patterns being of the major diagnostic value. Environmental disruptions of circadian rhythms, including shift work, travel across time zones, and irregular social schedules, tend to precipitate or exacerbate mood-related symptoms of mania or the manic episodes of BD [44]. Seasonal alternations are similarly linked to the occurrence of mania, manic episodes being more frequent in spring and summer. In contrast, depressive episodes peak in early winter and are less frequent in summer [45]. Neither pharmacological nor genetic animal models of mania include similar associations. In the CURD protocol, normal rhythms were disturbed by nine interventions, including day-night rhythms (Fig. 1b), sleep rhythms (Fig. 1c), attention and activity rhythms (Fig. 1d and 1e), the temperature regulation (Fig. 1f), light and noise interference (Fig. 1g and 1h), and security threat (Fig. 1i).

Beside the manic-like behaviours, CURD treated mice also had some biological and functional abnormalities. As shown in Fig. 5, the glymphatic system is malfunctional in CURD and CUMR group (Fig. 5a). The *c-fos* as a broad indicator of neuronal activity was increased in CURD and AMP treated mice (Fig. 5c), while the expression of 5-HTT in cortex were significantly reduced in CURD and AMP groups, being associated with the elevated extracellular 5-HT (Fig. 5f and 5g). The depression-related CUMR or COR regimens demonstrated the opposite changes in *c-fos*, 5-HTT and 5-HT compared with CURD and AMP groups (Fig. 5c-5g). Some of the changes outlined above were reported in mania patients. For example, the level of 5-HT metabolite 5-hydroxyindoleacetic acid (5-HIAA) is elevated in cerebrospinal fluid (CSF) in 14 hospitalised drug-free manic patients [46, 47].

Mania and the mania episode of BD are serious mental disorders. The shortage of the appropriate animal models severely hinders the advances in research of pathogenesis and pharmacological management of mania. We designed the novel mania model induced solely by environmental disturbances, which are easy to replicate. This model, we believe, will aid research into molecular pathways of mania and the pharmacological mechanisms of anti-mania drugs.

## Acknowledgments

This work was supported by the National Natural Science Foundation of China, BL [grant number 81871852]; Shenyang Science and Technology Innovation Talents Project, BL [grant number RC210251]; LiaoNing Revitalisation Talents Program, BL [grant number XLYC1807137], the Scientific Research Foundation for Returned Scholars of Education Ministry of China, BL [grant number 20151098], LiaoNing Thousands Talents Program, BL [grant number 202078], ‘ChunHui’ Program of Education Ministry, BL [grant number 2020703].

## Author contributions

A.V. and B.L. designed and supervised the study; B.C., M.J., W.G., D.Z., X.L. and M.X. collected the data in vivo and analysed the relevant data; B.C., S.W., Y.F., X.W. and L.C. performed the behavioural experiments in vivo and analysed the data; B.L. and A.V. wrote the manuscript.

## Data availability

Source data underlying the main and supplementary figures are available in Supplementary Data 1. The uncropped blots are available in Supplementary Information file. The data that support the findings of this study are available from the corresponding author Baoman Li upon reasonable request.

